# Household cockroaches carry CTX-M-15, OXA-48 and NDM-1, and share beta-lactam resistance determinants with humans

**DOI:** 10.1101/357079

**Authors:** Noah Obeng-Nkrumah, Appiah-Korang Labi, Harriet Blankson, Georgina Awuah-Mensah, Daniel Oduro-Mensah, Judelove Anum, James Teye, Solomon Dzidzornu Kwashie, Evariste Bako, Patrick Ferdinand Ayeh-Kumi, Richard Asmah

**Author notes:** These authors contributed equally to this work (Noah Obeng-Nkrumah, Appiah-Korang Labi).

## Abstract

**Aim:** Household insect pests, including cockroaches, have gained consideration as potential vectors for multidrug resistant pathogens of public health concern. This study was designed to investigate whether household cockroaches share beta-lactam resistance determinants with human inhabitants.

**Methods:** From February through July 2016, 400 cockroaches were systematically collected from 100 households. Whole insect homogenates and faecal samples from inhabitants of all included households were cultured for cephalosporin-resistant enterobacteria (CRe). The CRe were examined for AmpC, ESBL, and carbapenemase genes; antibiotic susceptibility patterns; and conjugative transfer of antibiotic resistance mechanisms. Clonal relationships between isolates were determined by multi-locus sequence typing (MLST).

**Results:** Twenty CRe were recovered from whole cockroach homogenates of 15 households. Five harbored ESBL genes (*2 bla*_CTX-M-15/TEM-1_; 1 *bla*_CTX-M-15/TEM-4_; 1 *bla*_TEM-24_; 1 *bla*_SHV-4_), and 3 carried carbapenemase genes (2 *bla*_NDM-1_ genes and 1 *bla*_OXA-48_ gene) all of which were transferrable by conjugation to *E. coli* J53 recipients. There was high clonal diversity with low inter-species similarity regardless of the beta-lactamase gene sequence. From 6 households, the pair of cockroach and human CRe shared the same antibiogram, ST and/or conjugable *bla*_ESBL_ gene sequence (house 34, *E. coli* ST9-*bla*_TEM-4_; house 37, *E. coli* ST44-*bla*_CTX-15/TEM-4_; house 41, *E. coli* ST443-*bla*_CTX-15/TEM-1_; house 49, *K. pneumoniae* ST231-*bla*_SHV-13_).

**Conclusion:** The findings highlight household cockroaches as reservoirs of CTX-M-15, OXA-48 and NDM-1 genes that share beta-lactam resistance determinants with humans.

## Introduction

The production of extended-spectrum beta-lactamases (ESBL), Class C cephalosporinases and carbapenemases constitute the primary antibiotic resistance mechanism in Enterobacteriaceae [1,2]. Together, these beta-lactamases confer resistance to all available beta-lactam antibiotics, and are associated with significantly high levels of co-resistance to other classes of broad spectrum antimicrobials [1–6]. Increasingly, the CTX-M-15 type ESBLs are becoming predominant in Ghana [7,8]. Occurrence of the recently described OXA-48 type carbapenemase and widespread reports of *bla_NDM-1_* genes across Africa further compound the outlook of the antibiotic resistance problem [9–13]. Indeed, emergence of such resistant bacteria is often related to misuse and abuse of beta-lactam antibiotics [14,15]. However, the burden of such pathogens is also hugely aided by poor sanitation [16–18]. Cockroaches, which widely colonize the environment, including human dwellings, may well act as vectors of antibiotic-resistant bacteria.

Cockroaches are abundant in Ghanaian households and are known to harbor an array of pathogens [19]. They often reside in household sewage pipe systems, which are a repository of diverse infectious microorganisms. That cockroaches carry pathogenic bacteria is well documented in literature [19–23]. Some of the pathogenic bacteria may be carriers of drug-resistance determinants, particularly beta-lactamases. However, only few studies have investigated AmpC, ESBL or carbapenemase resistance elements in bacteria from cockroaches[24]. Consequently, the vector potential of cockroaches in the dissemination of such multidrug-resistance mechanisms is largely under-reported. In this study, we report the presence of CTX-M-15-, OXA-48- and NDM-1- producing bacteria in household cockroaches with particular focus on the conjugability of beta-lactam resistance determinants and clonal transmission to humans.

## Methods

### Study design and setting

Between February and July 2016, a cross-sectional study was conducted to collect cockroaches and human faecal samples from households in Ashaley Botwe, an urban municipality setting in Accra, Ghana. The municipality has a population of approximately 78,215, with most households occupied by an average of 5.6 persons[25]. The major source of water is pipe borne, and most of the households have proper sewage disposal systems with flush toilets and standardized septic tanks. Hundred households were selected by systematic random sampling using the Kish method[26] which statistically allowed for equal chances of selecting any household in the community. The selected households were approximately 150 metres from each other. From each household, live indoor cockroaches were collected. Stool samples were also collected from inhabitants of each household. The study received ethical approval from the Ethics and Protocol Review Committee of the School of Biomedical and Allied Health Sciences, University of Ghana, with approval identification number: SBAHS-MD./10512194/aa/5a/2016-2017. All human participants provided written informed consents.

### Sample collection and processing

All 100 households were provided with 100 ml sterile containers with screw caps, sterile zipper bags containing gloves and forceps to use for capturing cockroaches. A member of each household was selected and educated appropriately to be responsible for capturing cockroaches. Only live cockroaches found indoors were captured for this study. Four cockroaches (irrespective of species) were requested per household and pooled as one sample. Each cockroach sample was soaked in 40 mL of Brain Heart Infusion broth (Sigma, UK), vortexed vigorously for 6 minutes, and ground with a sterile rod. The triturate was once again vortexed vigorously for 6 minutes to obtain a whole insect homogenate. A loop-full (approximately 10 μL) of suspension from each sample was inoculated onto SSI agar plate (SSI, Diagnostica, Denmark) with 30 μg cefpodoxime disk (MAST, UK) and incubated overnight at 37 °C. For stool samples, 1 gram of each specimen was suspended in 10 ml of sterile 0.9% physiological saline and vigorously vortexed. 1 ml of the suspension was cultured on SSI agar plates with 30 μg cefpodoxime disk. From each culture plate, distinct morphological phenotypes of enterobacteria growing around the cefpodoxime disks within an inhibition zone of 21mm were defined as screen positive for third generation cephalosporin resistance (CRe). Each distinct morphological phenotype was identified to the species level using the biochemical test kits Minibact-E^®^ (SSI, Diagnostica, Denmark) according to the manufacturer’s guidelines. Subsequently, four colonies of each speciated phenotype were subjected to the Kirby-Bauer method of susceptibility testing per guidelines of the Clinical and Laboratory Standard Institute (CLSI) using cefotaxime (30 μg) and ceftazidime (30 μg) antibiotics [27]. Isolates resistant to cefotaxime or ceftazidime were confirmed as CRe.

### Susceptibility test and assays for ESBL, AmpC and carbapenemase

All CRe isolates were tested for susceptibility to the following antibiotics (MAST, UK) according to CLSI guidelines[27]: ampicillin (10 μg), augmentin (10 μg/260 μg); meropenem (10 μg), tetracycline (30 μg), chloramphenicol (30 μg), cotrimoxazole (100 μg/240 μg), gentamicin (10 μg), ciprofloxacin (5 μg), nitrofurantoin (100 μg); piperacillin/tazobactam (10/30 μg), cefoxitin (30 μg), ceftaroline (30 μg), and tigecycline (30 μg). *Klebsiella pneumoniae* ATCC 700603 and *Escherichia coli* ATCC 25922 were used as quality control strains. Ceftaroline breakpoint was interpreted with European Committee for Antibiotic Susceptibility Testing (EUCAST) guidelines[28].Phenotypic detection of ESBL production was by the combination disk method with cefotaxime (30 μg) or ceftazidime (30 μg), alone or in combination with clavulanic acid (10 μg) [29]. AmpC expression was suspected in isolates with reduced susceptibility (inhibition zone <20 mm) to cefoxitin (30 μg). AmpC confirmation was by the use of cefotaxime (30 μg) or ceftazidime (30 μg) tablets, with or without boronic acid (250 ug) as per the manufacturer’s guidelines (Rosco Diagnostica, Taastrup, Denmark). An increase of ≥5 mm in zone diameter, due to boronic acid, was considered AmpC positive. Isolates with inhibition zone of <21 mm for 10 μg meropenem were considered carbapenem resistant [29]. Carbapenem resistant isolates were confirmed for class A and B carbapenemase using boronic acid (600 μg) and EDTA (750 μg) effect, respectively, on meropenem (10 μg). Strains with boronic acid or EDTA effect of ≥5 mm increase in zone diameter were considered positive for class A or B carbapenemase phenotype, respectively. Carbapenem resistant strains that were not susceptible to temocillin (30 μg) were considered positive for OXA-48 like carbapenemase. Carbapenem resistant strains were also subjected to the Modified Hodges Test [29].

### Genotypic characterization

Gene amplification and sequencing was done for ESBL-, AmpC-, and carbapenemase-producing isolates as well as their transconjugants. For each isolate, 10 μL of pure culture on Mueller Hinton agar was suspended in 300 μL Milli-Q ^®^ water, heated for 10 minutes at 98°C, and subsequently centrifuged for 5 minutes at 4°C and 20.000 g. The supernatant DNA lysate was transferred into sterile 1.5 ml Eppendorf^®^ tubes for storage at −5°C. See supplementary Table for amplification primers and conditions. For ESBLs, PCR was performed for *bla*_TEM_, *bla*_SHV_, *bla*_CTX-M-1_, *bla*_CTX-M-2_, *bla*_CTX-M-9_, *bla*_OXA-2_, *bla*_0XA-10_. Isolates with AmpC phenotypes were examined for 6 families of plasmid-mediated AmpC genes including MOX, CMY, DHA, ACC, EBC and FOX using the multiplex assay[30]. For carbapenemases, PCRs were designed for specific genes belonging to class A, B, and D carbapenemase phenotypes. A multiplex PCR assay was performed to differentiate 5 genes (GES, KPC, SME, INI-NMC-A) for class A carbapenemase, and 6 genes (IMP, VIM, GIM, SPM, SIM, and NDM-1) for class B phenotypes[31]. Isolates with Class D carbapenemase phenotypes were examined for OXA-48 like genes. Additional internal primers (supplementary data 1) were used for sequencing CTX-M-1, CTX-M-9, SHV and TEM genes. Nucleotide and deduced protein sequences were compared with sequences in the NCBI database (http://www.ncbi.nlm.nih.gov/BLAST). TEM and SHV beta-lactamase sequences were compared to wild-type *E. coli* TEM-1 and SHV-1 at http://www.lahey.org/studies. For Multilocus Sequence Typing (MLST), we used the the previously described *E. coli* protocol by Wirth et al. [32] and the PubMLST scheme for *K. pneumoniae* (http://pubmlst.org/kpneumoniae/). Seven housekeeping genes were amplified and sequenced for each *E.coli* (adk, fumC, gyr, icd, mdh, purA and recA) and *K. pneumoniae* (gapA, infB, mdh, pgi, phoE, rpoB, tonB). Resulting sequences of the seven house-keeping genes were analysed using the CodonCode Aligner software version 8.1 (Germany). The MLST sequence types (ST) and clonal complexes (CC) were assigned in accordance to the online platform PubMLST database. Phylogenetic minimum spanning tree was constructed using the online programme PHYLOViZ [33].

### Conjugation

All CRe isolates were included in the conjugation assay. Conjugations were done using sodium azide-resistant *E. coli* J53 as recipient [34]. None of the CRe showed resistance to sodium azide. The donors were cultured in Luria-Bertani (LB) (MAST, UK) broth with cefotaxime (8 μg/ml) overnight. The recipient was also cultured overnight in LB broth but with no antibiotic. Subsequently, 1 mL aliquots of each donor and the recipient were separately transferred into fresh 10 ml LB broth and incubated for 2 hours at 37°C. For each donor, 100 μL of culture was mixed with an equal volume of the recipient and the mixture was incubated for 6 hours at 37°C. Selection for transconjugants was carried out on MacConkey agar supplemented with sodium azide (150 mg/L) and cefotaxime (8 mg/L) or cefoxitin (32 mg/L) or meropenem (2 mg/L). Transconjugants were confirmed for ESBL, AmpC or carbapenemase genotype as previously described.

### Data analysis

Data were entered into a Microsoft Excel sheet for editing and analysis. Results are presented using descriptive analysis with proportions and percentages. Multidrug-resistant (MDR) isolates were those resistant to at ≥ 1 agent in ≥ 3 ntimicrobial categories (aminoglycosides, flouoroquinolones, penicillins, penicillins/β-lactamase inhibitors, antipseudomonal penicillins/β-lactamase inhibitors, cephamycins, anti-MRSA cephalosporins, 1^st^ and 2^nd^ generation cephalosporins, 3^rd^ and 4^th^ generation cephalosporins, monobactams, Carbapenems, polymixins, phosphonic acids, folate-pathway inhibitors, Glycylcyclines, phenicols). Extensively drug-resistant (XDR) isolates were non-susceptible to ≥ 1 agent in all but ≤ 2 antimicrobial categories. Conjugation frequency per recipient was expressed by dividing the number of transconjugants by the initial number of recipients.

## RESULTS

The study procedures and outcomes are summarized in Fig 1. All 100 insect homogenates yielded polymicrobial cultures after inoculation onto SSI agar with 10μg cefpodoxime disks. Fifteen of the homogenate cultures were screen positive for CRe. Of these, 10 grew only one dominating colony type and 5 had two morphologicaly diferent dominant isolates with distinct antibiograms corresponding to 20 screen positive Enterobacteriaceae isolates (Table 1). The 20 isolates were each resistant to cefotaxime or ceftazidime antibiotic by the Kirby-Bauer method of sensitivity testing, and were thus confirmed as cockroach CRe. Overall, 61 fecal samples were collected from all inhabitants of 15 households that had cockroach samples positive for CRe. The average number of inhabitants per household was 4±1.3. Fifteen (24.5%) of the 61 faecal samples were screen positive for CRe on SSI agar plate. These included 10 faecal cultures that yielded only one dominant colony type and 5 that cultured two morphologicaly different dominating isolates with completely different antibiograms corresponding to 20 screen positive CRe. All 20 screen positive CRe were resistant to cefotaxime or ceftazidime antibiotic disk, and were assigned human CRe.

**Fig 1.**
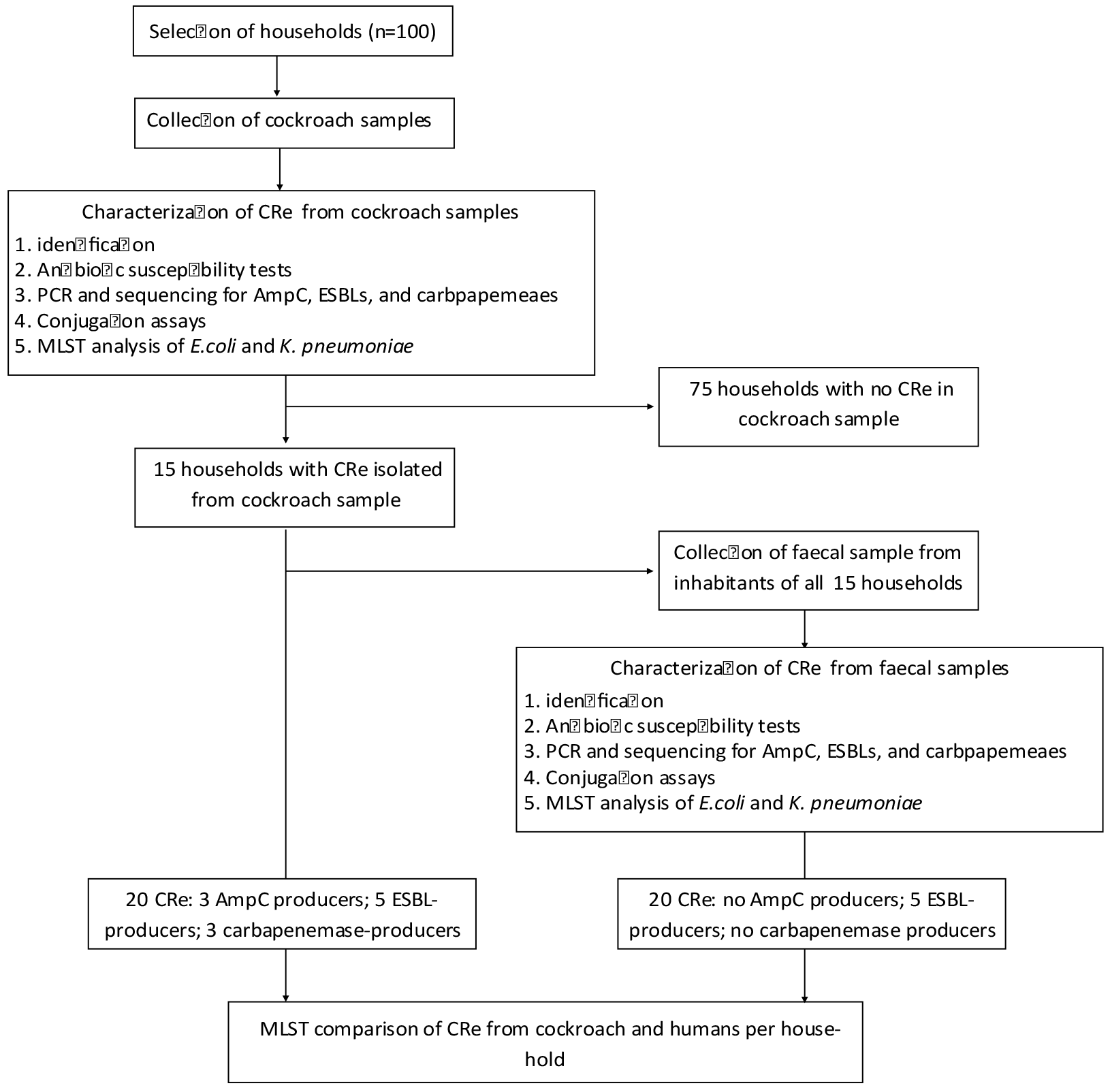
Summary of study protocols and outcomes. This is flow chart outline of study procedures and results.

### Characterization of CRe

Of the 20 cockroach CRe, 12 isolates (from 10 samples) expressed ESBL (n=5), AmpC (n=4) or were resistant to meropenem (n=3) (Table 1). The AmpC-producers were *Enterobacter freundii* (n=2) and *Enterobacter agglomerans* (n=2). Isolates with ESBL phenotype included 4 *E. coli* and 1 *K. pneumoniae.* Two of the 3 meropenem resistant isolates (*E. freundii* and *C. cloacae*) were found to be metallo-beta-lactamse (MBL)-producers. The third (*E. coli*) was non-susceptible to temocillin (30 ug); and was deemed presumptively positive for OXA-48 like carbapenemase (Table 1). None of the CRe isolates was positive for any combination of the three phenotypes. Of the 5 ESBL-producers, PCR and nucleotide sequencing identified *bla*_CTX-M-15/TEM-1_ in 2 *E. coli* isolates (Table 1). The remaining 3 harboured either *bla*_CTX-M-15/TEM-4_, *bla*_TEM-24_ or *bla*_SHV-3_ ESBL genes. None of the AmpC-producers had an identifiable plasmid-mediated AmpC gene. The 2 MBL-positive isolates each carried *bla*_NDM-1_ gene. The meropenem resistant *E. coli* with non-susceptibility to temocillin carried *bla*_OXA-48_ gene. Antibiogram of the 20 cockroach CRe revealed differences in resistance patterns between ESBL-, AmpC- or carbapenemase-positive isolates and those CRe negative for the 3 phenotypes (Table 1). The CRe without any of the phenotypes were MDR phenotypes in 3 of 8 isolates. They were susceptible to most of the non-beta-lactam antibiotics, with all 8 isolates susceptible to gentamicin (n=7/7), tigecycline (n=7/7), chloramphenicol (n=8/8), and nitrofurantoin (n=7/7). In contrast, MDR phenotype was reported in all isolates with ESBL or AmpC phenotype. Two of the 3 carbapenemase-positive CRe were also XDR. Table 2 shows the molecular characterization and antibiogram of all 20 human CRe. None was positive for AmpC or carbapenemase phenotype. Five CRe expressed ESBLs (4 *E. coli* and 1 *K. pneumoniae*). When the ESBL-producing faecal isolates were subjected to PCR and sequencing, the *K. pneumoniae* carried *bla*_SHV-3_. The 4 *E. coli* separately harboured *bla*_TEM-24_, *bla*_TEM-14_, *bla*_CTX-M-15/TEM-4_, and *bla*_CTX-M-15/TEM-1_. The antibiotic susceptibility profile of all 20 human CRe mirrored that observed for cockroach isolates with clear differences in resistance pattern between ESBL- and non-ESBL producers (Table 2).

**Table 1.**
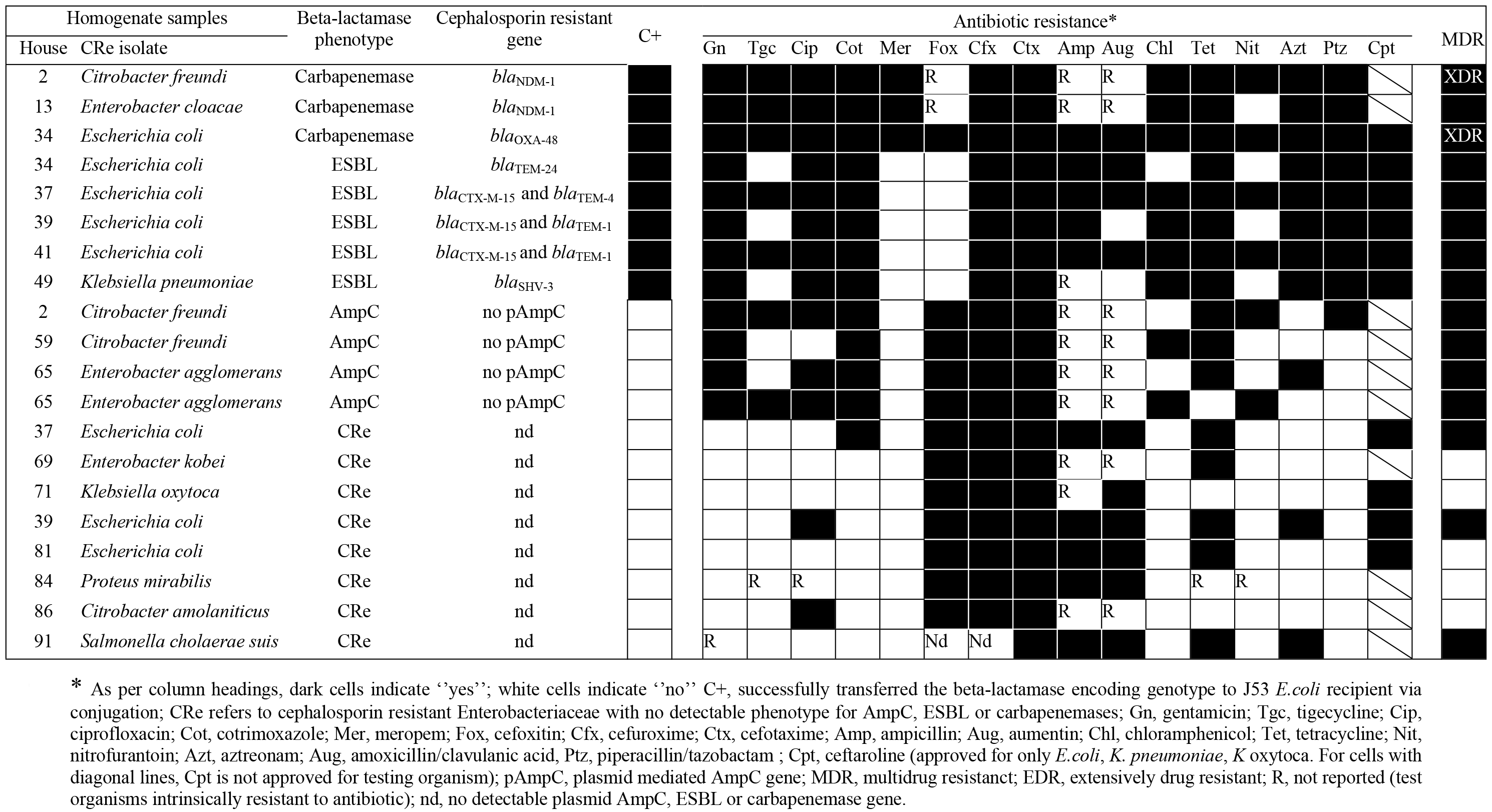
Characteristics of 20 CRe recovered from 100 whole insect homogenates.

**Table 2.** Characteristics of 20 CRe recovered from human inhabitants in households with Cockroach CRe

| House | CRE isolate | β-lactamase phenotype | Number of persons | Antibiotic resistance* |  |  |  |  |  |  |  |  |  |  |  |  |  |  |  | MDR | β-lactamase encoding genes | C+ |
| --- | --- | --- | --- | --- | --- | --- | --- | --- | --- | --- | --- | --- | --- | --- | --- | --- | --- | --- | --- | --- | --- | --- |
|  |  |  |  | Gn | Tgc | Cip | Cot | Mer | Fox | Cfx | Ctx | Amp | Aug | Chl | Tet | Nit | Azt | Ptz | Cpt |  |  |  |
| 2 | *Escherichia coli* | CRE | 1 of 3 |  |  |  |  |  |  |  |  |  |  |  |  |  |  |  |  |  | none found |  |
| 13 | *K. oxytoca* | CRE | 2 of 4 |  |  |  |  |  |  |  |  | R |  |  |  |  |  |  |  |  | none found |  |
| 13 | *Escherichia coli* | CRE |  |  |  |  |  |  |  |  |  |  |  |  |  |  |  |  |  |  | none found |  |
| 34 | *Escherichia coli* | ESBL | 1 of 5 |  |  |  |  |  |  |  |  |  |  |  |  |  |  |  |  |  | *bla*<sub>TEM-24</sub> |  |
| 37 | *Escherichia coli* | ESBL | 1 of 3 |  |  |  |  |  |  |  |  |  |  |  |  |  |  |  |  |  | *bla*<sub>CTX-M-15</sub> and *bla*<sub>TEM-4</sub> |  |
| 39 | *Escherichia coli* | CRE | 1 of 5 |  |  |  |  |  |  |  |  |  |  |  |  |  |  |  |  |  | none found |  |
| 41 | *Escherichia coli* | ESBL | 2 of 5 |  |  |  |  |  |  |  |  |  |  |  |  |  |  |  |  |  | *bla*<sub>CTX-M-14</sub> |  |
| 41 | *Escherichia coli* | ESBL |  |  |  |  |  |  |  |  |  |  |  |  |  |  |  |  |  |  | *bla*<sub>CTX-M-15</sub> and *bla*<sub>TEM-1</sub> |  |
| 49 | *Klebsiella pneumoniae* | ESBL | 1 of 5 |  |  |  |  |  |  |  |  | R |  |  |  |  |  |  |  |  | *bla*<sub>SHV-3</sub> |  |
| 59 | *Escherichia coli* | CRE | 1 of 3 |  |  |  |  |  |  |  |  |  |  |  |  |  |  |  |  |  | none found |  |
| 65 | *Escherichia coli* | CRE | 1 of 2 |  |  |  |  |  |  |  |  |  |  |  |  |  |  |  |  |  | none found |  |
| 69 | *Escherichia coli* | CRE | 2 of 3 |  |  |  |  |  |  |  |  |  |  |  |  |  |  |  |  |  | none found |  |
| 69 | *Klebsiella pneumoniae* | CRE |  |  |  |  |  |  |  |  |  |  |  |  |  |  |  |  |  |  | none found |  |
| 71 | *Escherichia coli* | CRE | 2 of 2 |  |  |  |  |  |  |  |  |  |  |  |  |  |  |  |  |  | none found |  |
| 71 | *Klebsiella pneumoniae* | CRE |  |  |  |  |  |  |  |  |  | R |  |  |  |  |  |  |  |  | none found |  |
| 81 | *Escherichia coli* | CRE | 2 of 5 |  |  |  |  |  |  |  |  |  |  |  |  |  |  |  |  |  | none found |  |
| 81 | *Citrobacter koseri* | CRE |  |  |  |  |  |  | R |  |  | R | R |  |  |  |  |  |  |  | none found |  |
| 84 | *Escherichia coli* | CRE | 1 of 3 |  |  |  |  |  |  |  |  |  |  |  |  |  |  |  |  |  | none found |  |
| 86 | *Escherichia coli* | CRE | 1 of 6 |  |  |  |  |  |  |  |  |  |  |  |  |  |  |  |  |  | none found |  |
| 91 | *Escherichia coli* | CRE | 1 of 6 |  |  |  |  |  |  |  |  |  |  |  |  |  |  |  |  |  | none found |  |

### Conjugation Assays

The 20 CRe each from cockroach and human inhabitants were subjected to conjugation experiments (Table 3). All CRe negative for ESBLs, AmpCs or carbapenemases could not transfer their cephalosporin resistance to *E. coli* J53 recipients. Similarly, none of the AmpC-producing isolates transferred the phenotype. Successful conjugative experiments were demonstrated only in ESBL- or carbapenemase-producing isolates (Table 3). For these isolates, PCR amplification and sequencing of the ESBL or carbapenemase genes in the *E. coli* J53 transconjugants showed the same *bla* gene types previously identified in the donor isolates. The ESBL genes in human isolates transferred at significantly lower conjugation frequencies (range: 1.1×10^−5^ - 1.9×10^−4^) compared to those from cockroach isolates (range: 2.3×10^−3^ - 4.8×10^−2^). For cockroach isolates, carbapenemase genes appeared to conjugate with lower efficiency (range: 1.1×10^−3^ - 1.9×10^−3^) compared to ESBL genes (range: 2.3×10^−3^ - 4.8×10^−2^). Resistance to non-β-lactam antimicrobials was also cotransferred in some cases, in addition to transfer of ESBL or carbapenemase phenotype. The most frequently co-transferred antibiotic resistance phenotype in transconjugants were cotrimoxazole, followed by tetracycline and then ciprofloxacin. Tigecycline and nitrofurantoin resistance did not transfer to recipients despite repeated attempts.

**Table 3.** Conjugation characteristics of ESBL- and carbapenemase-producing CRe isolated from cockroaches and humans

| House | Isolates | Donor isolates characteristics |  | *E. coli* J53 transconjugant |  |  |
| --- | --- | --- | --- | --- | --- | --- |
|  |  | β-lactamase type | Antibiotic resistant profile | β-lactamase type | Conjugation frequency | Cotransferred antibiotic resistance |
| 2 | *Citrobacter freundii*<sup>cockroach</sup> | Carbapenemase positive phenotype; *bla*<sub>NDM-1</sub>; | Gn, **Tgc**, Cip, Cot, Mer, Cfx, Ctx, **Chl**, Tet, **Nit**, Azt, Ptz | Carbapenemase phenotype; *bla*<sub>NDM-1</sub> | 1.1x10<sup>-3</sup> | Gn, Cip, Cot, Mer, Cfx, Ctx, Tet, Azt, Ptz |
| 13 | *Enterobacter cloacae*<sup>cockroach</sup> | Carbapenemase positive phenotype; *bla*<sub>NDM-1</sub> | Gn, **Tgc**, Cip, Cot, Mer, Cfx, Ctx, Chl, Tet, Azt | Carbapenemase type; *bla*<sub>NDM-1</sub> | 1.7x10<sup>-3</sup> | Gn, Cip, Cot, Mer, Cfx, Ctx, Chl, Tet, Azt, Ptz |
| 34 | *Escherichia coli*<sup>cockroach</sup> | Carbapenemase positive phenotype; *bla*<sub>OXA-48</sub> | Gn, **Tgc**, Cip, Cot, Mer, Fox, Cfx, Ctx, Amp, Aug, Chl, Tet, **Nit**, Azt, **Ptz**, Cpt | Carbapenemase phenotype; *bla*<sub>OXA-48</sub> | 1.9x10<sup>-3</sup> | Gn, Cip, Mer, Fox, Cfx, Ctx, Amp, Aug, Chl, Tet, Azt, Cpt |
| 34 | *Escherichia coli*<sup>cockroach</sup> | ESBL phenotype, *bla*<sub>TEM-24</sub> | Gn, Cip, Cot, Cfx, Ctx, Amp, Aug, Tet, Azt, **Ptz**, Cpt | ESBL phenotype, *bla*<sub>TEM-24</sub> | 2.3x10<sup>-2</sup> | Gn, Cip, Cot, Cfx, Ctx, Amp, Aug, Tet, Azt, Cpt |
| 37 | *Escherichia coli*<sup>cockroach</sup> | ESBL phenotype, *bla*<sub>CTX-M-15</sub> and *bla*<sub>TEM-4</sub> | **Gn**, **Tgc**, Cip, Cot, Cfx, Ctx, Amp, Aug, **Chl**, Tet, **Nit**, Azt, Ptz, Cpt | ESBL phenotype, *bla*<sub>CTX-M-15</sub> and *bla*<sub>TEM-4</sub> | 3.4x10<sup>-2</sup> | Cip, Cot, Cfx, Ctx, Amp, Aug, Tet, Azt, Ptz, Cpt |
| 39 | *Escherichia coli*<sup>cockroach</sup> | ESBL phenotype, *bla*<sub>CTX-M-15</sub> and *bla*<sub>TEM-1</sub> | **Gn**, **Cip**, Cot, Cfx, Ctx, Amp, **Chl**, Tet, Azt, Ptz, Cpt | ESBL phenotype, *bla*<sub>CTX-M-15</sub> and *bla*<sub>TEM-1</sub> | 5.1x10<sup>-2</sup> | Cot, Cfx, Ctx, Amp, Tet, Azt, Ptz, Cpt |
| 41 | *Escherichia coli*<sup>cockroach</sup> | ESBL phenotype, *bla*<sub>CTX-M-15</sub> and *bla*<sub>TEM-1</sub> | **Gn**, **Tgc**, **Cip**, Cot, Cfx, Ctx, Amp, Aug, **Chl**, Tet, **Nit**, Azt, **Ptz**, Cpt | ESBL phenotype, *bla*<sub>CTX-M-15</sub> and *bla*<sub>TEM-1</sub> | 3.9x10<sup>-2</sup> | Cot, Cfx, Ctx, Amp, Aug, Tet, Azt, Cpt |
| 49 | *Klebsiella pneumoniae*<sup>cockroach</sup> | ESBL phenotype, *bla*<sub>SHV-3</sub> | **Gn**, Cip, Cot, Cfx, Ctx, **Chl**, Tet, Azt, Ptz, Cpt | ESBL phenotype, *bla*<sub>SHV-3</sub> | 4.8x10<sup>-3</sup> | Cip, Cot, Crx, Ctx, Amp, Tet, Azt, Ptz, Cpt |
| 34 | *Escherichia coli*<sup>human</sup> | ESBL phenotype, *bla*<sub>TEM-24</sub> | Gn, Cip, Cot, Cfx, Ctx, Amp, Aug, Tet, Azt, **Ptz**, Cpt | ESBL phenotype, *bla*<sub>TEM-24</sub> | 1.6x10<sup>-6</sup> | Gn, Cip, Cot, Cfx, Ctx, Amp, Aug, Tet, Azt, Cpt |
| 37 | *Escherichia coli*<sup>human</sup> | ESBL phenotype, *bla*<sub>CTX-M-15</sub> and *bla*<sub>TEM-4</sub> | **Gn**, **Tgc**, Cip, Cot, Cfx, Ctx, Amp, Aug, **Chl**, Tet, **Nit**, Azt, Ptz, Cpt | ESBL phenotype, *bla*<sub>CTX-M-15</sub> and *bla*<sub>TEM-4</sub> | 2.1x10<sup>-5</sup> | Cip, Cot, Cfx, Ctx, Amp, Aug, Tet, Azt, Ptz, Cpt |
| 41 | *Escherichia coli*<sup>human</sup> | ESBL phenotype, *bla*<sub>CTX-M-15</sub> and *bla*<sub>TEM-1</sub> | **Gn**, **Tgc**, **Cip**, Cot, Cfx, Ctx, Amp, Aug, **Chl**, **Tet**, **Nit**, Azt, **Ptz**, Cpt | ESBL phenotype, *bla*<sub>CTX-M-15</sub> and *bla*<sub>TEM-1</sub> | 7.1x10<sup>-4</sup> | Cot, Cfx, Ctx, Amp, Azt, Cpt |
| 41 | *Escherichia coli*<sup>human</sup> | ESBL phenotype, *bla*<sub>CTX-M-14</sub> | **Gn**, **Tgc**, **Cip**, Cot, Cfx, Ctx, Amp, **Chl**, Tet, Azt, **Ptz**, Cpt | ESBL phenotype, *bla*<sub>CTX-M-14</sub> | 2.1x10<sup>-5</sup> | Cot, Cfx, Ctx, Amp, Aug, Tet, Azt, Cpt |
| 49 | *Klebsiella pneumoniae*<sup>human</sup> | ESBL phenotype, *bla*<sub>SHV-3</sub> | **Gn**, Cip, Cot, Cfx, Ctx, **Chl**, Tet, Azt, Ptz, Cpt | ESBL phenotype, *bla*<sub>SHV-3</sub> | 1.98x10<sup>-5</sup> | Cip, Cot, Crx, Ctx, Tet, Azt, Ptz, Cpt |

### Phylogenetic analysis

Overall, there were 20 Cockroach CRe and 20 human CRe. Of these, we compared by MLST the top 2 predominant isolates *E. coli* (n=23) and *K. pneumoniae* (n=4). There was high clonal diversity with low inter-species similarity regardless of the beta-lactamase gene sequence (Fig 2). Four *E. coli* STs and clonal complexes were found in cockroach CRe (ST48/CC10, ST101/CC101, ST367/CC23, ST405/CC405). Among human CRe, we found 5 *E. coli* STs with associated clonal complexes (ST88/CC23, ST162/CC469, ST189/CC165, ST215/CC10, ST341/CC205) and 5 singleton STs (ST117, ST542, ST871, ST1250, ST1287). The *K. pneumoniae* STs identified in only humans were ST171 and ST244 singletons. The following STs with clonal complex were detected in CRe from both human and cockroach samlples (*E. coli:* ST9/CC20, ST44/CC10, ST443/CC205, ST453/CC86; *K. pneumoniae:* ST231/CC86). The *E. coli* ST215 was identified in human CRe from 2 households. In one clonal complex (CC10), the genotype was detected in 3 households, all from human CRe.

**Fig 2.**
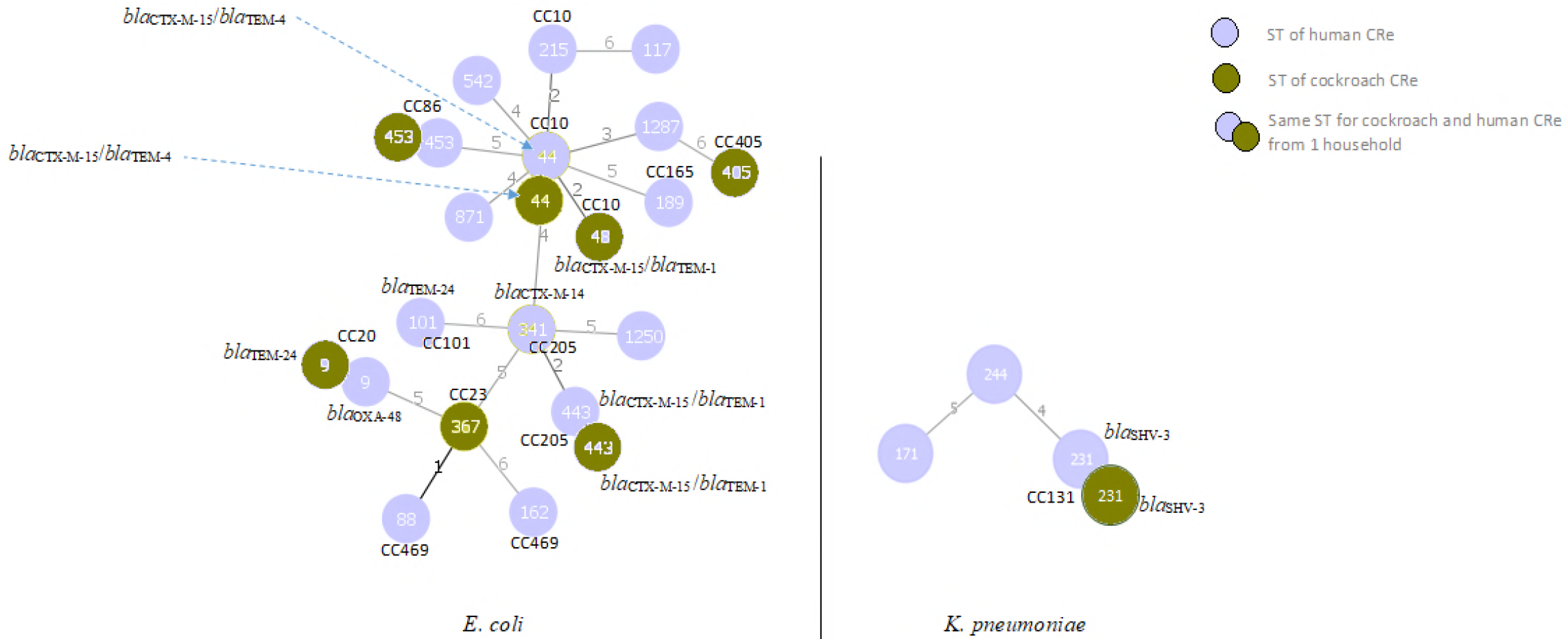
Minimum spanning tree based on MLST allelic profiles of CRe from human and cockroach samples. Minimum spanning tree based on MLST allelic profiles of cephalosporin resistant *E. coli* and *K. pneumoniae* in human and cockroach samples from household. Each circle represents an identified MLST sequence type (ST). The circle in red (or the lower half) represents an ST identified in cockroach CRe. The circle in blue (or the upper half) represents an ST identified in human CRe. The numbers on the connecting lines illustrate the number of differing alleles. The clonal complexes (CC) if present are indicated for the STs. Nd refers to CRe isolates without any detectable pAmpC, ESBL or carbapenemase gene.

### Comparison of cockroach and human CRe

In this study, all cockroach CRe isolates belonged to households with human inhabitants colonized by CRe. Thus there were 15 households with both cockroach and human CRe. Six households had human and cockroach CRe of the same bacteria identity (*E. coli* or *K. pneumoniae*) (Table 5). From 4 households, the pair of human and cockroach isolates shared the same ST and *bla*_ESBL_ gene sequence (house 34, *E. coli* ST9-*bla*_TEM-4_; house 37, *E. coli* ST44-*bla*_CTX-15/TEM-4_; house 41, *E. coli* ST443-*bla*_CTX-15/TEM-1_; house 49, *K. pneumoniae* ST231-*bla*_SHV-13_). The pair also had the same antibiogram; and successfully transferred their ESBL phenotype and genotype by conjugation to *E. coli* J53 recipients. In house 81, the CRe (from human and cockroach samples) belonged to the same *E. coli* ST453 but exhibited different antibiogram with no identifiable ESBL, AmpC or carbapenemase gene. The pair did not transfer their cephalosporin resistance phenotype by conjugation to *E. coli* J53 recipients. One human inhabitant (house 41) was colonized by *E. coli* ST341-*bla*_CTX-M-14_ which was different from the corresponding cockroach isolate (*E. coli* ST443 with *bla*_CTX-15/TEM-1_ ESBL gene). Both isolates however belonged to the same clonal complex CC205.

**Table 4.**
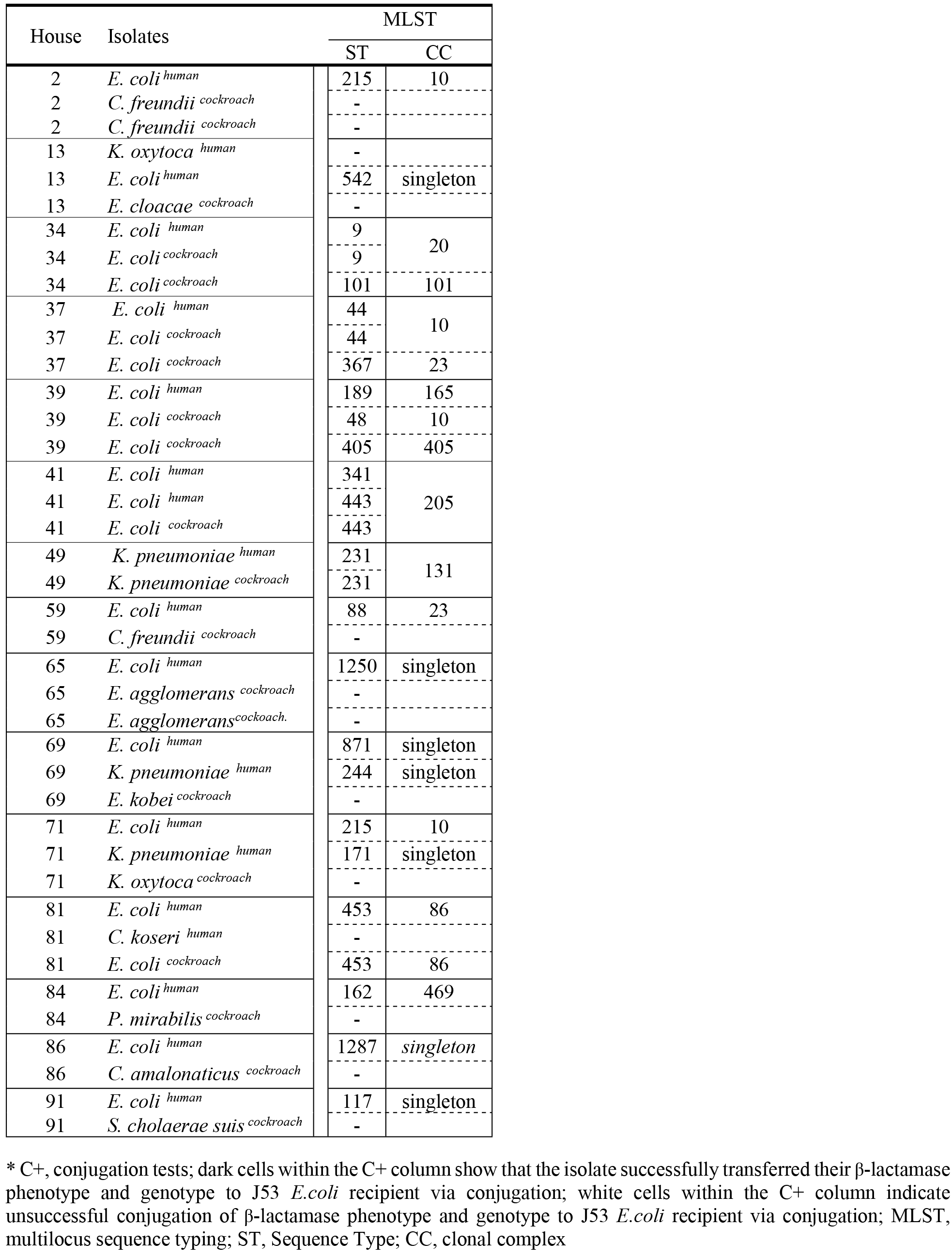
MLST analysis of CRe from cockroach and human inhabitants

## Discussion

The direct transfer of AmpC-, ESBL-, or carbapenemase-producing bacteria to humans via insects seems possible, but evidence is sparse [24,35]. Loucif et al. showed that 9 of 12 bacterial isolates from cockroaches in an Algerian hospital harboured CTX-M-15 genes. Of these, one expressed an OXA-48 type carbapenemase and belonged to the sequence type 528 [24]. Elsewhere, Fotedar and colleagues indicated in an experimental study that insects fed with *Pseudomonas aeruginosa* could excrete the bacteria even up to 114 days post-infection[35]. In our study, we report that cockroaches carry MDR CTX-M-15, XDR OXA-48, and XDR NDM-1 producing enterobacteria; and share the same STs and beta-lactam resistance determinants with humans in the household. There is paucity of data on community-based molecular studies examining cockroach colonization with carbapenemase producing Enterobacteriaceae, limiting the scope for comparing our results with existing literature. To our knowledge this is the first published description of *bla*_NDM-1_ and *bla*_OXA-48_ genes in enterobacteria recovered from household cockroaches.

Cockroaches often reside in household sewage systems. As they scurry, they contaminate surfaces including utensils leading to the horizontal transmission of bacteria from person to person. Our findings highlight the importance of cockroaches as potential reservoir of epidemiologically significant multidrug resistant pathogens of public health concern. Some observations merit attention. First, merely 4% of households had cockroach and human isolates from one house that shared the same ST or clonal complex, antibiogram, *bla*_ESBL_ gene sequence, and successfully transferred their ESBL phenotype and genotype by conjugation to *E. coli* J53 recipients. The results point to zoonotic transmission by clonal spread and highlight domestic cockroaches as constituting a ‘revolving door’ through which ESBL-producing enterobacteria disseminate to human contacts. The successful inter-genus conjugation events reported in this study suggest that, at least for the non-*E. coli* isolates used as donors, the plasmids involved are likely to be conjugative plasmids with broad host range. This hints at the potential for these resistance determinants to spread freely across bacterial species in the environment. Our results are strengthened by other findings that have reported direct transfer of ESBL-producing bacteria to humans via close contact with animal sources. Huijbers et al. revealed an increased risk for people with close broiler contact as well as strong indications of the transmission from animals[36]. In a comparative analysis of ESBL-Positive E. *coli* from animals and humans from the UK, the Netherlands and Germany, Wu et al. showed that about 1.2% of the animal isolates shared the same MLST CC with the human ones indicating that zoonotic transmission [37]. A direct transfer from poultry to people with close contact has also been reported [38]. Second, the finding of enterobacteria with *bla*_NDM-1_ and *bla*_OXA-48_ in household cockroaches is alarming given that the strains could very quickly disseminate, and originate uncontrollable dissemination of pandemic clones for which effective antibiotics may not be available in Ghana. The ease of transmission of *bla*_NDM-1_ genes have already been discussed by Karen Bush[2,39–41]. Interestingly, although the *bla*_NDM-1_ and *bla*_OXA-48_ genes were conjugable, no household inhabitant was faecally colonized with a carbapenem resistant isolate. In Ghana, there are no published data on carbapenemase, but it is the experience that carbapenem resistance is low[6,42,43]. Last, the antibiogram of isolates described in this study do not differ from that generally reported for clinical strains. In this study, the general antibiogram of the study isolates showed an overall high resistance prevalence to many routinely used non-beta-lactam antibiotics including ciprofloxacin and gentamicin. It is the experience in Ghana that resistance to these antibiotics among clinical Enterobacteriaceae is high[42,44,45]. It is widely presumed that such high levels of resistance are associated with plasmid-mediated beta-lactamases. Indeed, our results show a clear separation, in which, the prevalence of resistance was higher among strains expressing ESBL, AmpC or carbapenemase than those negative for the enzymes. The CRe isolates without ESBL, AmpC or carbapenemase were susceptible to several antibiotics tested. None of these CRe also successfully transferred their cephalosporin resistance phenotype to *E. coli* J53 recipients by conjugation, suggesting an intrinsic resistance borne on the chromosome. The results suggest that, from the clinical point of view, if we can succeed at curtailing the spread of ESBL-, AmpC-, or carbapenemase producing enterobacteria, we may somewhat succeed at reducing the prevalence of antimicrobial resistance among enterobacteria.

Our data should be interpreted considering potential limitations. Households were provided with cockroach collection kits and procedural guidelines. We are however unable to ascertain if participants strictly adhered to aseptic collection guidelines. Nonetheless, the fact that all the CRe isolates were the dominant colonies on culture suggest stable colonization. We have no data on antibiotic use by household contacts and are thus unable to relate CRe colonization and antibiotic consumption. Due to small sample size and provincial household concentration, our study is not meant to be representative for the prevalence of AmpC-, ESBL-, or carbapenemase-producing bacteria. It is noteworthy that the complimentary sampling of humans is biased since only humans with corresponding positive household cockroach samples were examined. The study was designed to investigate whether, within individual households, cockroaches and humans share isolates of the same clone and antibiotic resistance determinants of public health concern.

## Conclusion

We report the disturbing colonization of household cockroaches with multidrug resistant CTX-M-15 ESBL-producers, and extensively drug resistant NDM-1 and OXA-48 carbapenemase-positive *Enterobacteriacaea.* The findings highlight cockroaches as insects of public health concern; and calls for regulations on their control especially in healthcare settings.

## Acknowledgements

The authors are grateful to Bill Wickstead (of Wickstead Lab, University of Nottingham School of Life Sciences, UK) who assisted with some reagents for the project. We thank the following for their laboratory support: Michael Olu-Taiwo (University of Ghana School of Biomedical and Allied Health Sciences, Department of Medical Laboratory Sciences); Mary Magdalene Osei (University of Ghana School of Biomedical and Allied Health Sciences, Medical Microbiology department); Mr Thomas Dankwa (all of Korle-Bu Teaching Hospital Central Laboratory); and Mr Seth Agyeman (Korle-Bu Teaching Hospital Central Laboratory).

## Supporting information

S1 Table. Oligonucleotides and PCR conditions used for amplification (internal primers included for sequencing)

